# Oocytes maintain ROS-free mitochondrial metabolism by suppressing complex I

**DOI:** 10.1101/2022.05.27.493392

**Authors:** Aida Rodríguez-Nuevo, Ariadna Torres-Sanchez, Juan M Duran, Cristian De Guirior, Maria Angeles Martínez-Zamora, Elvan Böke

## Abstract

Oocytes form before birth and remain viable for several decades before fertilisation. Although poor oocyte quality accounts for the majority of female fertility problems, little is known about how oocytes maintain cellular fitness, nor why they eventually decline with age. Reactive oxygen species (ROS) produced as by-products of mitochondrial activity are associated with lower rates of fertilisation and embryo survival. Yet, how healthy oocytes balance essential mitochondrial activity with the production of ROS is unknown. Here, we show that oocytes evade ROS by remodelling the mitochondrial electron transport chain (ETC) through elimination of complex I. Combining live-cell imaging and proteomics in human and *Xenopus* oocytes, we find that early oocytes exhibit greatly reduced levels of complex I. This is accompanied by a highly active mitochondrial unfolded protein response, which is indicative of an imbalanced ETC. Biochemical and functional assays confirm that complex I is neither assembled nor active in early oocytes. Thus, we report the first physiological cell type without complex I in animals. Our findings clarify why patients suffering from complex I related hereditary mitochondrial diseases do not experience subfertility, in contrast to diseases involving other mitochondrial complexes. Complex I suppression represents an evolutionary-conserved strategy that allows longevity while maintaining biological activity in long-lived oocytes.

## MAIN

Human primordial oocytes are formed during foetal development and remain arrested in the ovary for up to 50 years. Despite a long period of dormancy, oocytes retain the ability to give rise to a new organism upon fertilisation. Decline in oocyte fitness is a key contributor to infertility with age^1^. However, little is known about how oocytes maintain cellular fitness for decades to preserve their developmental potential, complicating efforts to understand the declining oocyte quality in ageing mothers.

Primordial oocytes remain metabolically active during dormancy^2,3^ and thus, must maintain mitochondrial activity for biosynthesis of essential biomolecules^4^. Yet, mitochondria are a major source of cellular ROS, which form as by-products of mitochondrial oxidative metabolism. Although ROS can function as signalling molecules^5^, at high concentrations, ROS promote DNA mutagenesis and are cytotoxic. Indeed, ROS levels are linked to apoptosis and reduced developmental competence in oocytes and embryos^6–8^. However, the mechanisms by which oocytes maintain this delicate balance between mitochondrial activity with ROS production have remain elusive.

### Mitochondrial ROS in early oocytes

Early human oocytes can only be accessed via invasive surgery into ovaries. Therefore, biochemical investigations into oocyte biology has historically been hindered by severe sample limitations. As a consequence, mitochondrial activity in primordial oocytes remains largely unstudied. Here, we overcome these challenges by utilising an improved human oocyte isolation protocol recently developed in our laboratory^2^, which we combine with a comparative evolutionary approach using more readily-available *Xenopus* stage I oocytes (both referred to as early oocytes from hereafter). This approach allowed us to generate hypotheses using multi-species or *Xenopus-only* analyses, and subsequently test those hypotheses in human oocytes.

We began our studies by imaging live early human and *Xenopus* oocytes labelled with various mitochondrial probes that quantify ROS levels. Neither *Xenopus* nor human early oocytes showed any detectable ROS signal, whereas mitochondria in somatic granulosa cells surrounding the oocytes displayed high ROS levels and served as a positive control (Fig1a-c; S1a-e).

**Figure 1.**
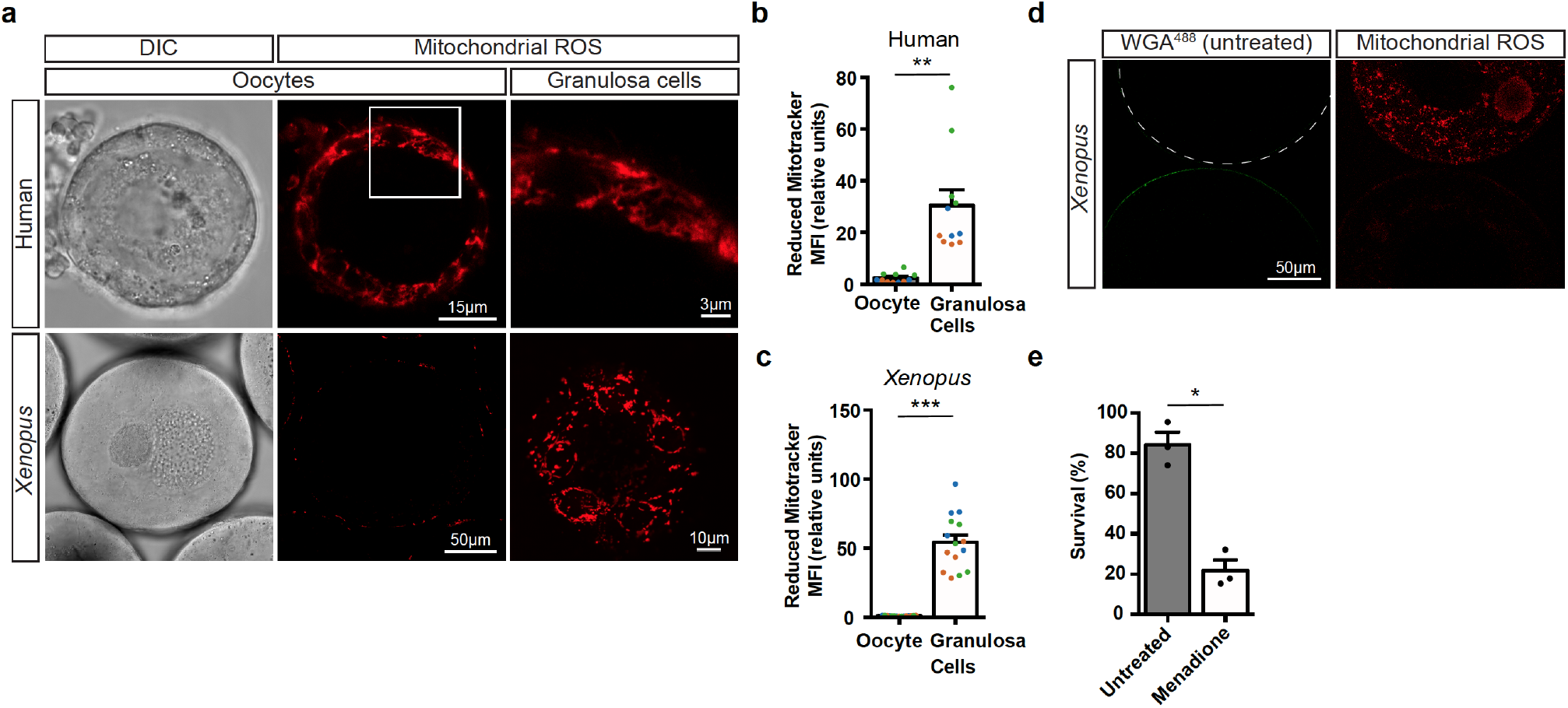
Early oocytes have undetectable levels of ROS. **a,** Live-cell imaging of human and *Xenopus* early oocytes with attached granulosa cells labelled with MitoTracker Red CM-H_2_XRos (H2X), a reduced mitochondrial dye that does not fluoresce until it is oxidized by ROS. *Xenopus* granulosa cells were imaged at the basal plane of the oocyte. **b, c,** Quantification of the mean fluorescence intensity (MFI) of H2X in the oocyte and in the population of granulosa cells surrounding the equatorial plane of the oocyte of human (**b**) and *Xenopus* (**c**). Data represent mean ± SEM. \*\**p* = 0.0001, \*\*\**p* = 4.13 × 10^-11^ n=3 biological replicates shown in 3 colours. **d,** Untreated and menadione treated (dashed line) *Xenopus* early oocytes were combined in the same dish and labelled with MitoSOX. Untreated oocytes were previously incubated with Wheat Germ Agglutinin (WGA)^488^ (green) to mark the plasma membrane. **e,** Overnight survival of early oocytes after menadione treatment. At least 50 early oocytes were incubated per condition. Data represent mean ± SEM, n=3. * *p* = 0.0016.

To distinguish between the possibilities that low ROS-probe levels resulted from low ROS production or alternatively, a high scavenging capacity to eliminate ROS, we treated *Xenopus* oocytes with menadione. Mild treatment with menadione promotes the formation of ROS by introducing electrons directly to complex III^9^ but does not affect survival negatively in cell lines and fruit flies^10,11^. After a mild treatment with menadione (10 μM for 2 hours), we could indeed detect ROS in oocyte mitochondria. However, the majority of oocytes (78.3%) could not recover and died when they were left to recover overnight (Fig S1f, Fig1d,e). These results suggest that evasion of ROS damage in oocytes involves a tight control of ROS generation, rather than a higher scavenging capacity of oocytes against ROS.

### Mitochondrial respiration in early oocytes

Mitochondrial ETC activity generates ROS, and establishes the mitochondrial membrane potential, which enables oxidative-phosphorylation-(OXPHOS)-coupled ATP synthesis^12^. Using dyes that sense membrane potential (TMRE and JC-1), we found that mitochondria in human and *Xenopus* early oocytes exhibit lower membrane potentials compared to neighbouring granulosa cells (Fig2a-d, S2a,b). Undetectable ROS levels and low membrane potential suggest that mitochondria in early oocytes have either low or absent ETC activity. To differentiate between these two possibilities, we investigated whether oocytes perform OXPHOS and measured respiration rate in *Xenopus* oocytes. Low ETC activity would cause a reduction in basal mitochondrial respiration, while in the absence of ETC activity, mitochondrial respiration would not be detectable. Granulosa cell-stripped early oocytes exhibited a low basal respiration rate but a similar maximal respiration rate compared to growing oocytes (Fig2e-g). This respiration was efficiently coupled to ATP synthesis, resulting in an undetectable proton leak (Fig2e). Therefore, we conclude that mitochondria in early oocytes have a functional ETC, with low activity.

**Figure 2.**
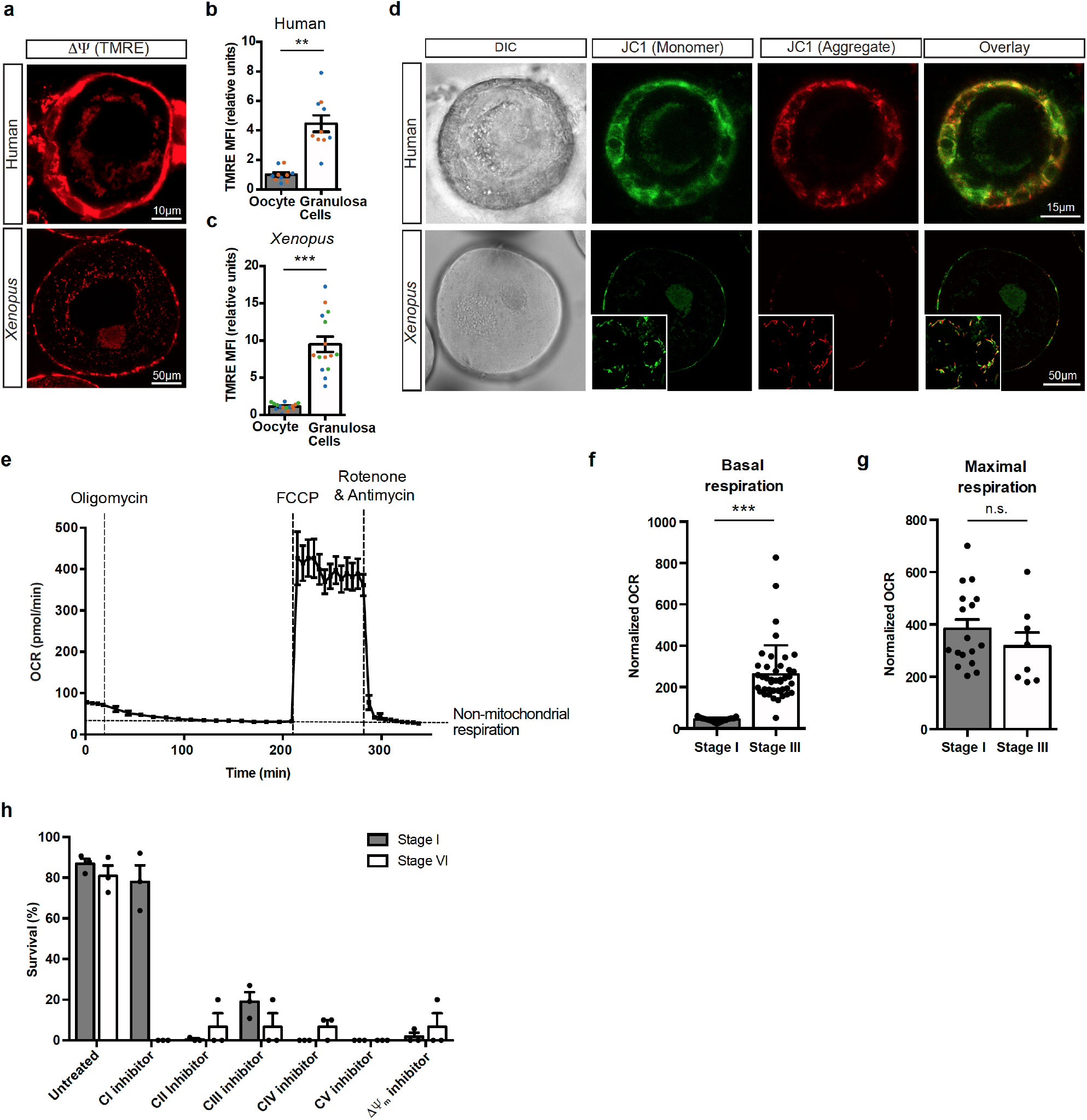
OXPHOS is low, but essential in early oocytes. Live-cell imaging of human and *Xenopus* early oocytes with attached granulosa cells labelled with **a,** Tetramethyl rhodamine (TMRE) to detect mitochondrial membrane potential (ΔΨ_m_); and **d,** JC-1, a membrane potential sensitive binary dye. Green JC-1 fluorescence is a sign of low membrane potential, while red fluorescence indicates JC-1 aggregation inside mitochondria, and thus, high membrane potential. Insets in *Xenopus* images represent granulosa cells imaged in the basal plane of the oocyte. Scale bars are as indicated in the figure. n ≥ 3; see figures S2a-b for JC-1 quantifications. **b, c,** Quantification of the mean fluorescence intensity (MFI) of TMRE inside the oocyte and in the population of granulosa cells surrounding the equatorial plane of the oocyte for human (**b**) and *Xenopus* oocytes (**c**). Data represent mean ± SEM. ** *p* = 1.21 × 10^-5^ n=2 in human and *** *p* = 8.91 × 10^-9^ n=3 in *Xenopus*; replicates shown in colours. **e,** Oxygen consumption rate (OCR) as assessed by a seahorse analyser (XFe96) in early (stage I) *Xenopus* oocytes (mean ± SEM, n = 9). **f,** Basal and **g**, maximal oxygen consumption rate in early (stage I) and growing (stage III) *Xenopus* oocytes, normalized for total protein/sample (mean ± SEM, n = 8). *** *p* = 2.98 × 10^-8^, n.s. not significant (*p* = 0.299) **h,** Overnight survival of early (stage I) and late (stage VI) oocytes after treatments with mitochondrial poisons: complex I (CI) to V (CV) inhibitors and an ionophore (5 μM Rotenone, 50 mM malonic acid, 5 μM antimycin A, 50 mM KCN, 200 μM DCCD or 30 μM CCCP, respectively). At least 50 early- and 10 late-stage oocytes were incubated per condition. ΔΨ_m_: mitochondrial membrane potential. Data represent mean ± SEM, n=3 biological replicates.

To assess the importance of individual complexes of the OXPHOS machinery (ETC complexes and ATP synthase [complex V]), for oocyte health, we performed a series of survival experiments with *Xenopus* oocytes. Exposing oocytes to inhibitors specific for each OXPHOS complex, we found that early and late-stage oocytes all died upon treatment with inhibitors of complexes II, III, IV and V (malonate, antimycin A, KCN, and DCCD, respectively) (Fig 2h, S2c). Although late-stage oocytes all died upon treatment with the complex I inhibitor rotenone, 78% of early oocytes survived in presence of rotenone (Fig 2h, S2c). The drastically reduced survival of oocytes by dissipation of membrane potential (by CCCP) or by individual inhibition of complexes II to V indicates that mitochondrial respiration is essential for oocyte survival, and that activities of complexes II to V are fundamental in early oocytes. The insensitivity of early oocytes to complex I inhibition indicates that they do not utilise complex I as an essential entry port for electrons and suggests that early oocytes rely on complex II for electron entry. However, we cannot rule out the possibility that complex I may still function in untreated oocytes.

### Quantitative proteomics of mitochondria in oocytes

So far, we have shown that mitochondria in early oocytes have an apparent lack of ROS, low membrane potential, low basal respiration rates and rotenone resistance in culture. We next wanted to investigate the mechanistic basis of this unusual physiological state of mitochondria in early oocytes.

Mitochondrial proteomic lists have been successfully used for hypothesis generation to understand mitochondrial pathways in different tissues, but a proteomic characterization of oocyte mitochondria has been missing. Therefore, we purified mitochondria from early and late-stage *Xenopus* oocytes, and performed MultiNotch proteomics including muscle mitochondria as a somatic cell control (Fig3a). 926 mitochondrial proteins were identified (and 807 quantified) in three biological replicates from wild type outbred animals, representing 80% of known mitochondrial proteins (Table 1, Fig S3a). Although mitochondrial proteome in diverse cell types could be quite different^13^, we found comparable levels of mitochondrial “house-keeping” proteins (such as import complex TIMMs and TOMMs) across different maturity stages (Fig S3b; Table 1) enabling us compare and contrast changes in other pathways.

**Figure 3.**
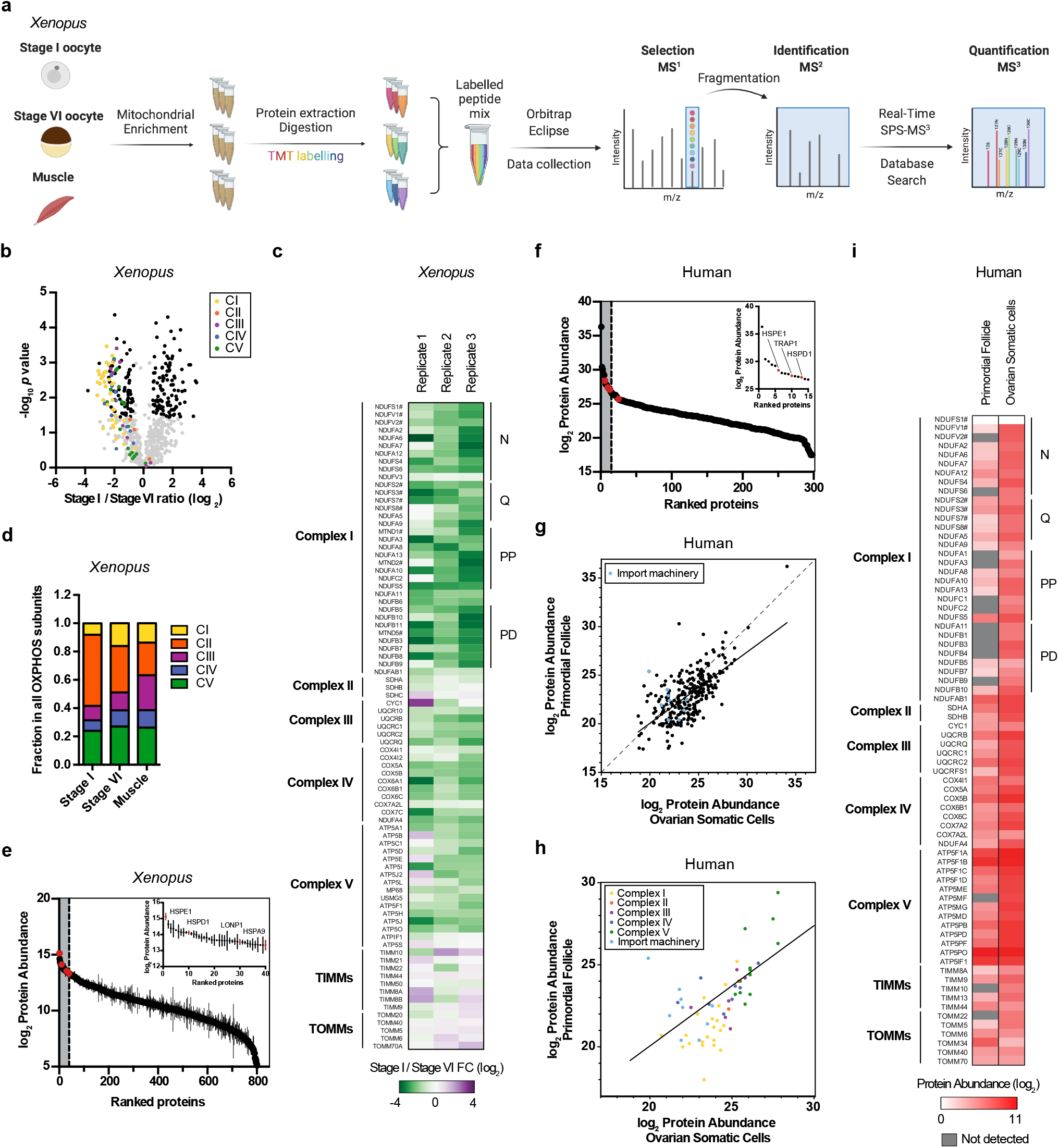
Mitochondrial proteomes of *Xenopus* and human oocytes. **a,** A schematic representation of quantitative proteomics experiment. **b,** Volcano plot showing *p* values versus fold changes of mitochondrial proteins between early (stage I) and late (stage VI) oocytes. Mitochondrial OXPHOS machinery are indicated in colours depicted in the figure. Other mitochondrial proteins significantly changing (q value <0.05, >1.5-fold change) are depicted in black. n=3 outbred animals. **c,** Heatmap of fold changes (early-vs late-stage oocytes) of all quantified subunits of the OXPHOS and mitochondrial import machinery. # marks core subunits of complex I. **d,** Fraction of mean abundance of the indicated complexes among all of OXPHOS machinery in oocytes and muscle tissue. See figure S3d for error bars. Note that this analysis inevitably misses undetected subunits. CI - CV: Complex I - Complex V **e,** Early *Xenopus* (stage I) oocytes proteome ranked by abundance. Inset represents top 5% most abundant proteins. UPR^mt^ proteins are indicated in red. Data are mean ± SEM from n=3. **f,** Human primordial follicle proteome ranked by abundance. Inset represents top 5% most abundant proteins. Oocytes were collected from ovaries of two patients and pooled together. UPR^mt^ proteins are indicated in red. **g,h,** Scatter plot comparing mitochondrial (**g**) and OXPHOS (**h**) protein abundance in human primordial follicles and ovarian somatic cells. Dashed line represents slope=1 and solid line is the linear regression model representing the relationship between mitochondrial proteomes of primordial follicles and ovarian somatic cells. **i,** Heatmap of normalized mitochondrial protein abundances in human primordial follicles and ovarian somatic cells organized by subunits of OXPHOS and mitochondrial import machinery. # marks core subunits of complex I. Proteins that were not identified are indicated in grey.

Most ETC subunits showed a lower absolute abundance in early oocytes compared to late-stage oocytes (Fig3b,c), and to muscle (FigS3c), which is expected due to the presence of fewer cristae in mitochondria of early oocytes^14–16^. The analysis of the relative abundance of each complex to total abundance of subunits of OXPHOS machinery in each cell type revealed low levels of complexes I, III and IV, and a significant enrichment of complex II in early oocytes (Fig3d; S3d). In support of our findings with the ETC inhibitors (Fig 2h, S2c), the depletion of complex I in early oocytes was the most pronounced from all other ETC complexes (Fig3b-d).

Furthermore, among the most abundant proteins in the early oocyte mitochondria were mitochondrial proteases and chaperones (Fig3e; S3e,f; S4a). These proteins are upregulated upon activation of mitochondrial unfolded protein response (UPR^mt^)^17–19^, which is often triggered by an imbalance of ETC subunits in mitochondria.

We next asked whether complex I subunits were also depleted in human oocytes. A recent improvement in oocyte isolation protocols^2^ enabled us to perform a proteomic characterisation of early (primordial) oocytes in humans. Early oocytes and ovarian somatic cells were isolated from patient ovarian cortices, and analysed by semi-quantitative proteomics, which provided the increased sensitivity essential for the limited biological material. We identified 454 mitochondrial proteins (Table 2) (298 and 397 proteins were quantified for early oocyte and somatic cell samples, respectively), representing 40% of all known mitochondrial proteins. Here too, levels of mitochondrial import proteins TIMMs and TOMMs were similar between oocytes and ovarian somatic cells (Fig3g-i) demonstrating an equivalent mitochondrial abundance that facilitated comparison of protein levels between different cell types. The upregulation of proteins related to UPR^mt^ was also conserved in human early oocytes, further confirmed with immunofluorescence (Fig3f; S4b). An analysis of the OXPHOS machinery comparing oocytes and ovarian somatic cells revealed that, in line with the *Xenopus* data, many complex I subunits were either at very low levels or not identified in human oocytes (Fig3h,i). A relative abundance analysis of ETC subunits again revealed a remodelling in early oocytes, with particularly low levels of complex I and high levels of complex V (FigS3g). Thus, we concluded that both the low abundance of complex I subunits and the imbalance of complex I to other complexes of the OXPHOS machinery are conserved between human and *Xenopus* early oocytes.

In conclusion, our proteomic characterisation of mitochondria revealed an overall reduction of ETC subunits in early oocytes of human and *Xenopus*, with complex I levels displaying the strongest disproportionate depletion.

### Complex I is not assembled in early oocytes

Taken together, our proteomics and survival experiments suggest the hypothesis that both early human and *Xenopus* oocytes remodel their ETC to decrease complex I levels to an extent that complex I becomes unnecessary for survival. This result is very surprising, because no animal cells have been shown to be able to dispense with complex I and only one other multicellular eukaryote, the parasitic mistletoe, is known to dispense with complex I entirely^20^. Therefore, we directly assayed complex I assembly status and function in early oocytes, using colorimetric, spectrophotometric, and metabolic assays.

We first investigated complex I assembly status in oocytes, which is tightly linked to its function^21^. Complex I is a multi-modular ~1 MDa complex composed of 14 core and 31 accessory subunits in humans, some of which are essential for its proper assembly and function. The individual absence of some of these subunits causes the loss of specific modules from the fully assembled complex, affecting the overall size and function of complex I^21^. To investigate whether complex I is fully assembled in early oocytes, we first examined our proteomics data to see whether there was any specific downregulation of a particular complex I module in early oocytes. Levels of subunits belonging to N, Q, PP and PD modules of complex I were compared between *Xenopus* early and late-stage oocytes. However, no significant changes were observed in terms of ratios among different modules in early oocytes (FigS5a).

The size of complex I in native protein gels has been used as a tool to reveal the assembly status of the complex^21,22^. Thus, we compared mitochondria isolated from early oocytes to those from late stage oocytes, and from muscle tissue of *Xenopus* and mice as somatic cell controls by blue-native (BN)-PAGE followed by complex I in-gel activity assays. Strikingly, complex I was neither fully-assembled nor active in early oocytes (Fig4a, S5b). BN-PAGE followed by an immunoblot against a complex I subunit, NDUFS1, confirmed that early oocytes do not have any detectable fully-assembled complex I (Fig S5c). Denaturing SDS-PAGE gels also verified comparable mitochondrial loading and very low protein levels of complex I subunits in early oocytes (Fig4b). To rule out any possibility of immunoblotting detection problems, areas corresponding to assembled complex I and complex II from BN-PAGE gels were analysed by proteomics and all theoretical complex I and II subunits are shown (Fig S5d, Table 3). Although complex II subunits were detected at comparable levels in all samples, the majority of complex I subunits were not detected in early oocytes (Fig4c). Thus, we conclude that complex I is not fully assembled in early oocytes.

**Figure 4.**
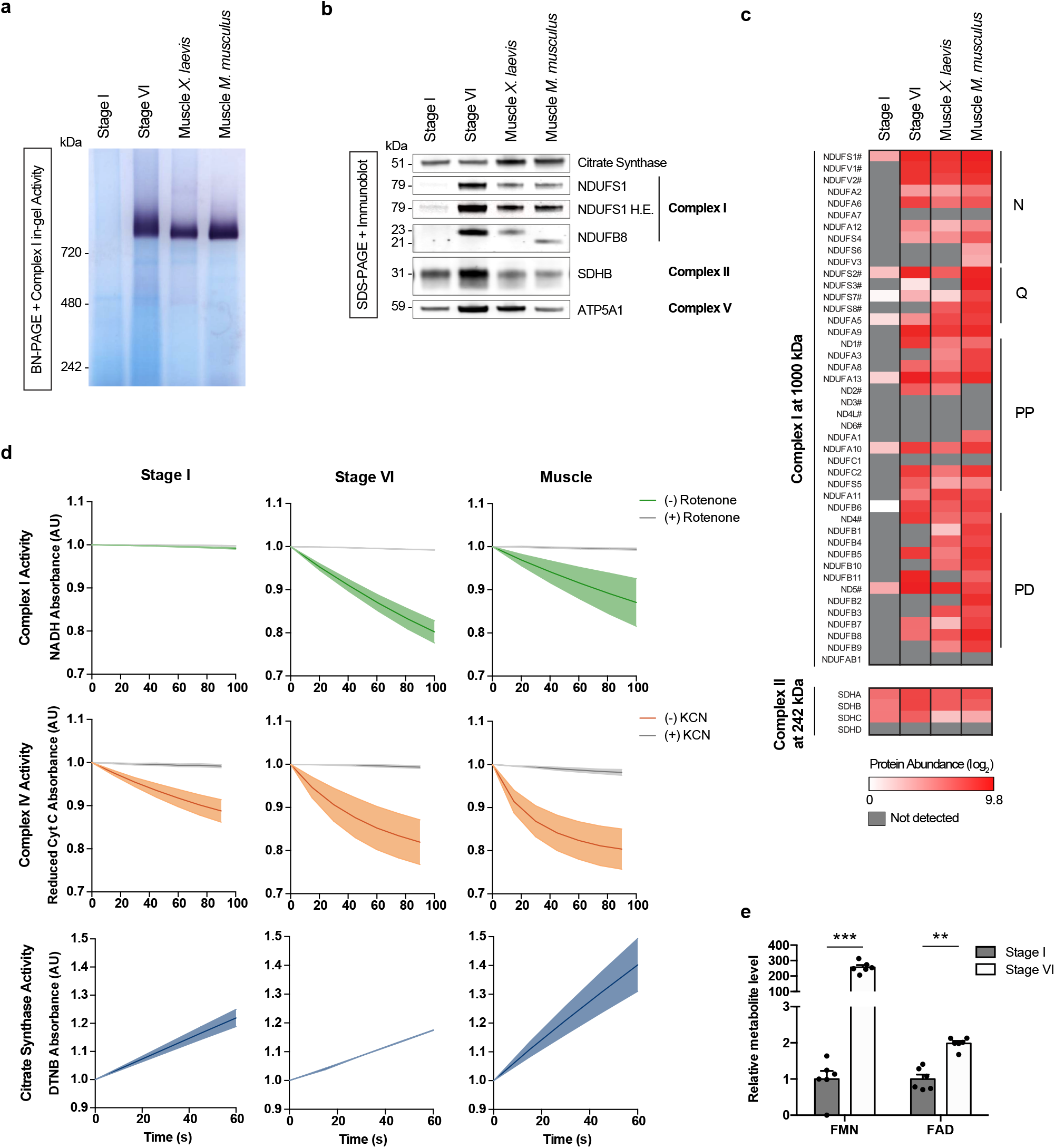
Complex I is not assembled in early oocytes. **a,** DDM solubilized mitochondrial fractions were resolved by BN-PAGE and complex I activity was assayed by NTB reduction in the presence of NADH. n ≥ 3, see figure S5b for quantifications. **b,** SDS-PAGE and immunoblotting of mitochondrial fractions in (**a**) for complex I (NDUFS1 and NDUFB8), complex II (SDHB) and complex V (ATP5A1) subunits, and citrate synthase as mitochondrial loading control. H.E. denotes high exposure. **c,** Heatmap of protein abundances detected in bands excised from regions corresponding to assembled complex I, and complex II. Complex I subunits are organized by module, and # marks core subunits. Proteins that were not quantified are indicated in grey. **d,** Spectrophotometric analysis of complex I (green, rotenone-specific activity), complex IV (orange, KCN-specific activity) and citrate synthase (blue) activities in mitochondrial extracts from early (stage I), late (stage VI) oocytes and muscle. Data represent mean ± SEM, n=3 biological replicates. **e,** FMN and FAD levels in early (stage I) and late (stage VI) oocytes. Data represent mean ± SEM, n=6. *** *p* = 6.92 × 10^-9^, ** *p* = 3.57 × 10^-5^.

Next, we checked for the presence of active subassemblies of complex I that might not be detected by in-gel activity assays. We examined NADH:CoQ oxidation in isolated mitochondrial membranes from early and late-stage oocytes, as well as muscle tissue to measure substrate consumption by complex I. We also measured Complex IV and citrate synthase activities to ensure presence of mitochondrial activity in these samples. Complex IV and citrate synthase activities were detected in all three samples. However, complex I activity was absent in early oocyte samples, in contrast to late stage oocyte samples and muscle samples.

Finally, to validate the absence of complex I in early oocytes by metabolomics, we checked the levels of flavin mononucleotide (FMN), an integral part of complex I in early and late stage oocytes. Although levels of another flavin nucleotide, FAD, were within a 2-fold range between these stages, FMN levels were about 200-fold higher in late-stage oocytes, compared to the low levels detected in early oocytes (Fig 4e). The remarkable depletion of FMN is complementary evidence supporting complex I deficiency in early oocytes.

Combining the *in vivo* evidence with proteomics and biochemical assays *in vitro*, our results demonstrate that early oocytes avoid ROS by eliminating one of the main ROS generators in the cell, mitochondrial complex I. Complex I subunits are reduced to such low levels that complex I cannot be fully assembled nor its activity be detected in early oocytes. This reveals a new strategy utilized by long-lived oocytes of *Xenopus* and humans to maintain a low-ROS mitochondrial metabolism. To our knowledge, early oocytes are the first and only physiological cell type in animals that exist without a functional mitochondrial complex I.

## Discussion

Oocytes are formed during female embryogenesis but perform their main function only after the onset of sexual maturation. Hence, these unique cell types, endowed with the transfer of an organism’s genetic material in pristine condition, must survive many years in the female body while minimizing damage to their cytoplasmic and nuclear components.

Here, we have shown that dormancy involves survival with an inactive mitochondrial complex I. Although they have active organelles and transcription, dormant oocytes do not grow or divide. Therefore, they have low energy needs, which could be served by glycolysis and the lower membrane potential established by electron flow from complex II. We can speculate that adjustment of mitochondrial respiration allows longevity while maintaining biological activity. By shutting down complex I and keeping the rest of the OXPHOS system active, early oocytes keep their mitochondria polarised to support the synthesis of heme, essential amino acids and nucleotides, while keeping their activity low to avoid ROS. Other quiescent cells, such as neuronal and hematopoietic stem cells exhibit similarly low ROS levels, and reduced ETC activity^5,23^, raising the possibility that this regulatory mechanism might be utilised by additional cell types. Furthermore, UPR^mt^ is activated in early oocytes (Fig 3e,f; S4), likely in response to an imbalance of ETC complexes caused by the absence of complex I. Given that UPR^mt^ activation itself is sufficient to increase the lifespan of *C.elegans* and mouse^17–19^, we speculate that complex I inhibition further enhances the longevity of oocytes via its downstream activation of UPR^mt^. The causal relationships between these interacting factors and oocyte lifespan remain as a fascinating future direction to investigate.

Moreover, the discovery of complex I inactivity in early oocytes explains why complex I related mitochondrial pathologies (such as Leber’s Hereditary Optic Neuropathy [LHON]) do not lead to subfertility nor selection against homoplasmic mtDNA mutations that occur in other types of ETC dysfunction^24–26^. Because oogenic mitochondrial bottleneck happens in early oogenesis^27^, there would not be a selective pressure against mutations affecting an inactive complex I.

Our findings reveal yet another unique aspect of physiology oocytes have evolved to balance their essential function of beginning life with the requirement for longevity. Could complex I deficiency in primordial oocytes be exploited for other purposes? Some cancers seen in young women are highly treatable, however, their treatment leads to a severe reduction of the ovarian reserve and reduced prospects of motherhood. Drugs against complex I exist, and are already proposed for cancer treatments^28^. Future studies will show whether repurposing complex I antagonists can improve chemotherapy-related infertility, and thus life quality of young female cancer survivors.

## Supporting information

Tabel 2

Tabel 3

## METHODS

### Ethics

Ethical Committee permission to work with primordial oocytes from human ovary samples was obtained from the Comité Étic d’Investigació Clínica CEIC-Parc de salut MAR (Barcelona) and Comité Ético de investigación clínica CEIC-Hospital Clínic de Barcelona with approval number *HCB/2018/0497*. Written informed consent was obtained from all participants prior to their inclusions in the study.

All animals mentioned in the manuscript were sacrificed by accredited animal facility personnel before their tissues were extracted.

### Animal Models

*Xenopus laevis* adult females were purchased from Nasco (NJ, USA) and maintained in water tanks in the following controlled conditions: 18-21°C, pH 6.8-7.5, O2 4-20 ppm, conductivity 500-1500 μs, ammonia < 0.1 ppm. The C57BL/6J mice used in the experiments were purchased from Charles River laboratories and maintained in the Animal Facility of the Barcelona Biomedical Research Park (PRBB, Barcelona, Spain, EU) under specific pathogen-free conditions at 22°C, in a 12 hours light/dark cycle, and with access to food and water *ad libitum*. Female mice of 7 weeks-of-age were used for extracting muscle tissue.

### Oocyte isolation and culture

#### Human primordial oocytes

Ovaries were provided by the gynaecology service of Hospital Clinic de Barcelona, from women aged 19 to 34 undergoing ovarian surgery and were processed as previously described^1^. Briefly, ovarian cortex samples were digested in DMEM containing 25 mM HEPES and 2 mg/mL collagenase type III (Worthington Biochemical Corporation, LS004183) for 2 hours at 37°C with occasional swirling. Individual cells were separated from tissue fragments by sedimentation and collagenase was neutralized by adding 10% FBS (Thermo, 10270106). Follicles were picked manually under a dissecting microscope. All human oocyte imaging experiments were conducted in DMEM/F12 medium (Thermo, 11330-032) containing 15 mM HEPES and 10% FBS (Thermo, 10270106).

#### *Xenopus* oocytes

Ovaries were dissected from young adult (aged 3 to 5 years) female *Xenopus laevis* that had undergone euthanasia by submersion in 15% Benzocaine for 15 minutes. Ovaries were digested using 3 mg/ml Collagenase IA (Sigma, C9891-1G) in MMR by gentle rocking until dissociated oocytes were visible, for 30 to 45 minutes. The resulting mix was passed through two sets of filter meshes (Spectra/Mesh, 146424, 146426). All washes were performed in MMR. For live-imaging experiments with intact granulosa cells, oocytes were transferred to oocyte culture medium (OCM)^2^ at this stage. For the rest of the experiments, oocytes were stripped of accompanying granulosa cells by treatment with 10 mg/ml trypsin in PBS for 1 minute, followed by washes in MMR. Removal of granulosa cells was confirmed by Hoechst staining of a small number of oocytes.

### HeLa Cell culture

HeLa cells were confirmed to be mycoplasma negative and grown in DMEM (Thermo, 41965039) supplemented with 1mM sodium pyruvate (Thermo, 11360070) and 10% FBS (Thermo, 102701060).

### Live-cell imaging

Human or *Xenopus* early oocytes were labelled in their respective culture media (see above). Human oocytes were imaged using a 63x water immersion objective (N.A1.20, Leica, 506346) with an incubation chamber maintained at 37°C and 5% CO_2_. Frog oocytes were imaged using a 40x water immersion objective (N.A 1.10, Leica, 506357) in OCM at room temperature and atmospheric air, unless stated otherwise. All images were acquired using a Leica TCS SP8 microscope with the Leica Application Suite X (LAS X) software. Mean fluorescence intensity in granulosa cells and oocytes were quantified using Fiji software.

#### ROS probes

Oocytes and associated granulosa cells were incubated in 500 nM MitoTracker Red CM-H2Xros (Thermo, M7513) for 30 minutes, 5 μM MitoSOX Red (Thermo, M36008) for 10 min, or in 5 μM CellROX for 30 minutes. Cells were then washed and imaged in 35 mm glass bottom MatTek dishes in culture media, except CellROX labelling, for which MMR was used for imaging to satisfy the manufacturer’s instructions.

#### Mitochondrial membrane potential probes

Oocytes and associated granulosa cells were labelled for 30 minutes in 500 nM Tetramethylrhodamine ethyl ester perchlorate (TMRE) (Thermo, T669), or 45 minutes in 4 μM JC-1 (Abcam, ab141387). Cells were then washed and imaged in 35 mm glass bottom MatTek dishes.

### Oxygen Consumption Rate

Oxygen consumption rate (OCR) of *Xenopus* oocytes was measured using a Seahorse XFe96 Analyser (Agilent). Granulosa cell-stripped oocytes were placed in XFe96 culture plates immediately after their isolation in Seahorse XF DMEM Medium pH 7.4 supplemented with 10 mM Glucose, 1 mM Pyruvate and 2 mM Glutamine (Agilent; 103015-100, 103577-100, 103578-100, 103579-100). Cartridge was loaded with concentrated inhibitor solution to achieve 5 μM oligomycin, 2 μM carbonyl cyanide 4- (trifluoromethoxy) phenylhydrazone (FCCP) or a combination of 0.5 μM rotenone and 0.5 μM antimycin A. Mock medium injections were performed to account for inhibitor-independent decline in OCR. For each sequential injection, at least 4 measurement cycles were acquired consisting of 20 seconds mix; 90 seconds wait and 3 minutes measure, in at least 3 replicates. For basal and maximal respiration rates, assay-independent OCR decline was corrected, and non-mitochondrial respiration (resistant to rotenone-antimycin mix) was subtracted. OCR measurements for growing oocytes (Stage III; with a diameter of 450-600 μm^3^) had to be performed statically because the probe of the Analyzer compressed and destroyed these large oocytes in long-term measurements. For growing (stage III) oocytes, OCR was acquired during 5 cycles per well, each cycle being 20 seconds mix; 90 seconds wait and 3 minutes measure, in at least 4 replicates. The well size imposed a technical limitation on the maximum amount of oocytes per well (100 early and 8 growing oocytes), thus, respiration data were normalized for the total protein amount per sample.

### Treatments with OXPHOS inhibitors

At least 50 early (stage I) and 10 late (stage VI) oocytes were placed in 35 mm glassbottom dishes (MatTek) and incubated for 16 hours at 18°C in OCM with or without the addition of different mitochondrial inhibitors: 5 μM Rotenone (Sigma, R8875), 50 mM malonic acid (Sigma, M1296), 5 μM antimycin A (Abcam, ab141904), 50 mM potassium cyanide (KCN) (Merck Millipore, 1049670100), 200 μM N,N-dicyclohexylcarbodiimide (DCCD) (Sigma, D80002) and 30 μM carbonyl cyanide m-chlorophenyl hydrazone (CCCP) (Abcam, ab141229). Survival was assessed by counting the number of oocytes before and after treatments with intact morphology. Cell death in stage VI oocytes was recognized by the development of a mottling pattern in the pigmentation of the animal pole^4^. Images were taken by a Leica IC90 E stereoscope.

At least 50 early (stage I) oocytes were treated with 10 μM menadione (Sigma, M5625) or left untreated, for 2 hours in OCM, and washed into fresh OCM. A small fraction of untreated oocytes was labelled with Wheat Germ Agglutinin (WGA)^488^ (Biotium, 29022-1) to mark their plasma membrane and mixed with a small fraction of menadione treated oocytes in a glass bottom MatTek dish 4 hours after menadione was removed. The mixed population of oocytes were then labelled with MitoSOX and imaged. Survival of menadione treated or untreated oocytes was assessed by counting the number of oocytes immediately after menadione treatment (t=0) and after 16 hours in fresh OCM.

### Mitochondrial-enriched extracts

Mitochondrial enriched fractions were obtain as described in Frezza et al. for gastrocnemius muscle and with minor adaptations for oocyte samples^5^. Freshly isolated early oocytes from *Xenopus* were lysed in mitochondria buffer (MB: 250 mM Sucrose, 3 mM EGTA, 10 mM Tris pH 7.4), and spun at low speed to remove debris. The resulting supernatant was centrifuged at 20.000 g for 20 minutes at 4°C. Late oocytes were spin-crashed, and yolk-free fraction was combined 1:1 with MB and centrifuged at 20.000 g for 20 minutes at 4°C to pellet mitochondria. Mitochondrial pellets from early and late-stage oocytes were resuspended in MB and subjected to DNase treatment for 10 minutes and proteinase K treatment for 20 minutes. PMSF was added to stop proteolytic activity and samples were centrifuged again at 20.000 g for 20 minutes at 4°C. Protein concentration was estimated and aliquots of crude mitochondria were stored at −80°C until use.

### Spectrometric assessment of enzymatic activities of mitochondrial complexes

Mitochondrial complex I, complex IV and citrate synthase specific activities were determined as described before with minor modifications^6^. Briefly, mitochondrial extracts were subjected to three freeze-thaw cycles in hypotonic buffer (10 mM TrisHCl) before activity analysis using an Infinite M200 plate reader (Tecan) in black bottom 96 well plates (Nunc) at 37°C. For complex I NADH:CoQ activity assessment, reaction solutions (50 mM KP pH 7.5, 3 mg/mL BSA, 300 μM KCN, 200 μM NADH) with or without rotenone (10 μM) were distributed into each well first. Mitochondrial extracts were then added and NADH absorbance at 340 nm was measured for 2 minutes to establish baseline activity. The reaction was then started by the addition of ubiquinone (60 μM). NADH absorbance was recorded for 15 minutes every 15 seconds.

For complex IV activity assessment, reaction solutions (50 mM KP pH 7, 60 μM reduced CytC) with or without KCN (600 μM) were distributed into each well first, and reduced cytochrome c absorbance at 550nm was recorded for 2 minutes to establish baseline oxidation. Mitochondrial extracts were then added and absorbance was measured for 15 minutes every 15 seconds.

For citrate synthase activity, reaction solution (100 μM Tris pH 8, 0.1% Triton x-100, 100 μM DTNB, 300 μM Acetyl CoA) was distributed into each well first. Mitochondrial extracts were then added and absorbance at 410 nm was measured for 2 minutes to set the baseline, then the reaction was started by addition of the substrate oxaloacetic acid (500 μM). Production of TNB (yellow) was recorded by measuring the absorbance at 410 nm for 15 minutes every 15 seconds. Enzymatic assays were plotted with the baseline represented as 1 for simplicity.

### Denaturing SDS gel electrophoresis

Oocytes were collected after isolation, frozen in liquid nitrogen and kept at −80°C until further use. Samples were processed as described in Gupta et al. 2018^7^. Gastrocnemius total homogenates were obtained as described^8^. HeLa cells were lysed in RIPA buffer (50 mM Tris-HCl, 150 mM NaCl, 1% Nonidet P-40, 01% SDS, 1 mM EDTA, supplemented with protease inhibitor cocktail [Complete Roche Mini, 1 tablet per 50 ml]) and spun at 20.000 g to eliminate cell debris. Cell lysates or mitochondrial enriched fractions were resolved by SDS-PAGE using 4-12% NuPAGE Bis-Tris gels.

### Blue-Native PAGE (BN-PAGE) electrophoresis, and in-gel activity assays

Mitochondrial content in samples of different cell types (different stages of oocytes and muscle tissue) was first assessed by western-blotting for their citrate synthase levels. Next, similar amounts of mitochondrial fractions were solubilized in 1% DDM, and were resolved in native state using NativePAGE 3-12% Bis-Tris (Thermo, BN1001BOX) gradient gels as described before^9^. Complex I activity in-gel assays were performed as described in Jha et al^10^. Briefly, immediately after the run, BN-PAGE gels were incubated in 2 mM Tris pH 7.4, 0.1 mg/mL NADH, 2.5 mg/mL NTB to asses NADH:FMN electron transfer, denoted by the appearance of dark purple colour. Reduced NTB intensities were normalized to citrate synthase levels of the same samples, detected by SDS-PAGE followed by immunoblotting. Intensity measurements were performed using Fiji software.

### Immunoblot analysis

Denaturing SDS-PAGE gels were transferred to nitrocellulose membranes via wet-transfer using a Mini Trans-Blot Cell (Bio-Rad). Membranes were blocked in Intercept (TBS) Blocking Buffer (LI-COR), and incubated overnight at 4°C with primary antibodies diluted in Intercept 0.05% Tween-20 as follows: anti-ATP5A1 (Abcam; ab14748), anti-Citrate synthase (Abcam; ab96600), anti-GAPDH (Thermo; AM4300), anti-HSPE1 (Thermo, PA5-30428), anti-NDUFB8 (Abcam; ab110242), anti-NDUFS1 (Abcam; ab169540), and anti-SDHB (Abcam; ab14714). Primary antibodies were washed with TBS-T (0.05% Tween-20) and membranes were incubated in secondary antibodies antimouse IgG, DyLight 680 (Thermo, 35518) or anti-rabbit IgG DyLight 800 4X PEG (Thermo, SA5-35571). After washing, membranes were imaged by a near-infrared imaging system (Odyssey LI-COR). Densitometric analysis of immunoblotting images was performed using Fiji software.

BN-PAGE gels were transferred to PVDF membranes using a Mini Trans-Blot Cell (Bio-Rad). After wet-transfer, PVDF membranes were distained in methanol, blocked and incubated with antibodies against NDUFS1 (Abcam, ab169540) and ATP5A1 (Abcam, ab14748) for complex I and complex V immunodetection, respectively.

### FMN metabolomics analysis

Metabolomics data was generated by Metabolon (Durham, NC). Samples were prepared using the automated MicroLab STAR®system from Hamilton Company in the presence of recovery standard for QC. After protein precipitation in methanol, metabolites were extracted and analysed by ultrahigh performance liquid chromatography-tandem mass spectrometry (UPLC-MS/MS) by negative ionization. Raw data was extracted, peak-identified and QC processed using Metabolon’s hardware and software.

### Immunostaining paraffin ovary sections

Human and frog ovaries were fixed in 4% PFA in PBS overnight at 4°C, washed, embedded in paraffin blocks and cut in 5 μM sections. After deparaffinization, antigen retrieval was performed by heating the slides 15 minutes in 10 mM sodium citrate pH 6. Sections were blocked and permeabilized in 3% BSA, 0.05% Tween 20, 0.05% Triton x-100 each for 1 hour at room temperature. Sections were incubated overnight at 4°C in the presence of primary antibodies (1:100): anti-ATP5A1 (Abcam; ab14748) and anti-HSPE1 (Thermo, PA5-30428); then 2 hours at room temperature with secondary antibodies (1:500). Antibodies and dyes used: Goat anti-rabbit Alexa488 or Alexa-555 (1:500, Invitrogen, A-11008, A-21428), goat anti-mouse Alexa647 (Invitrogen, A21236), and Hoechst dye (1:500, Abcam, ab145597). A droplet of mounting medium (Agilent, S302380) was added onto the section before imaging with a Leica TCS SP8 microscope equipped with 40x (N.A 1.30, Leica 506358) and 63x (N.A 1.40, Leica 506350) objectives.

### Statistical analysis

All data are expressed as mean ± SEM. A simple linear regression was performed to fit a model between the mitochondrial protein abundances of primordial follicle and ovarian somatic cells samples (Fig 3g,h). Unpaired two-tailed Student’s t-test was used in all other analysis, *p* values were specified in figure legends, and < 0.05 were considered significant. Multiple t-tests were used in figures 4e, S2a, S2b and were corrected by the Sidak-Bonferroni method using GraphPad Prism. For *Xenopus* mass spectrometry protein levels comparison, q-values were calculated as adjusted *p* values and significance was considered for q-value < 0.05. Fold change heatmap was represented using JMP (version 13.2) software.

### Mass Spectrometry

#### Sample preparation

Quantitative Proteomics for *Xenopus:* Mitochondrial extracts from early (stage I) oocyte, late (stage VI) oocyte and gastrocnemius muscle were quantified and 100μg of each sample was processed with slight modifications from ^7^. In brief, methanol precipitated proteins were dissolved in 6M Guanidine hydrochloride (GuaCl). Samples were then digested with LysC (20 ng/μl) in 2M GuaCl overnight at room temperature. The next morning, samples were further diluted to 0.5M GuaCl and digested with Trypsin(10 ng/μl) and additional LysC (20 ng/μl) for 8 hours at 37°C. Later, samples were speed-vacuumed, and the resulting pellet was resuspended in 200 mM EPPS pH 8.0. 10 μl of TMT stock solutions (20 μg/ μl in acetonitrile) were added to 50 μl of samples, and samples were incubated 3 hours at room temperature. TMT was quenched with 0.5 % final concentration of hydroxylamine. The samples were combined in one tube, acidified by 10% phosphoric acid, and subjected to a MacroSpin C18 solid-phase extraction (SPE) (The Nest Group, Inc) to desalt and isolate peptides. TMT mixes were fractionated using basic pH reversed-phase fractionation in an Agilent 1200 system. Fractions were desalted with a MicroSpin C18 column (The Nest Group, Inc) and dried by vacuum centrifugation^11^.

Label-free proteomics for human oocytes: Human primordial follicles and ovarian somatic cells were collected from two individuals who underwent ovarian surgery. Samples were dissolved in 6M Guanidine hydrochloride (GuaCl) pH 8.5, diluted to 2M GuaCl and digested with LysC (10 ng/μl) overnight. Samples were further diluted down to 0.5M GuaCl and digested with LysC (10 ng/μl) and Trypsin (5 ng/μl) for 8 hours at 37°C. Samples were acidified by 5% formic acid and de-salted with home-made C18 columns.

Detection of Complex I and II subunits from BN-PAGE gels: Gel bands were destained, reduced with dithiothreitol, alkylated with iodoacetamide, and dehydrated with acetonitrile for trypsin digestion. After digestion, peptide mix was acidified with formic acid prior to LC-MS/MS analysis.

#### Chromatographic and mass spectrometric analysis

TMT and label free samples were analysed using a Orbitrap Eclipse mass spectrometer (Thermo Fisher Scientific) coupled to an EASY-nLC 1200 (Thermo Fisher Scientific). Peptides were separated on a 50-cm C18 column (Thermo Fisher Scientific) with a gradient from 4% to 32% acetonitrile in 90 min. Data acquisition for TMT samples was done using a Real Time Search MS3 method (RTS-MS3)^12^. The scan sequence began with an MS1 spectrum in the Orbitrap. In each cycle of data-dependent acquisition analysis, following each survey scan, the most intense ions were selected for fragmentation. Fragment ion spectra were produced via collision-induced dissociation (CID) at normalized collision energy of 35% and they were acquired in the ion trap mass analyser. MS2 spectra were searched in real time with data acquisition using the “PHROG” database^13^ with added mitochondrially encoded proteins. Identified MS2 spectra triggered the submission of MS3 spectra that were collected using the multinotch MS3-based TMT method^14^.

Label free samples were acquired in data-dependent acquisition (DDA) and full MS were detected in the Orbitrap. In each cycle of data-dependent acquisition analysis, the most intense ions were selected for fragmentation. Fragment ion spectra were produced via high-energy collision dissociation (HCD) at normalized collision energy of 28% and they were acquired in the ion trap mass analyzer.

Gel bands were analysed using a LTQ-Orbitrap Velos Pro mass spectrometer (Thermo Fisher Scientific, San Jose, CA, USA) coupled to an EASY-nLC 1000 (Thermo Fisher Scientific). Peptides were separated on a 25-cm C18 column (Nikkyo Technos Co., Ltd. Japan) with a gradient from 7% to 35% acetonitrile in 60 min. The acquisition was performed in data-dependent acquisition (DDA) mode and full MS were detected in the Orbitrap. In each cycle, the top twenty most intense ions were selected for fragmentation. Fragment ion spectra were produced via collision-induced dissociation (CID) at normalized collision energy of 35% and they were acquired in the ion trap mass analyser. Digested bovine serum albumin was analysed between each sample and QCloud^15^ was used to control instrument performance.

#### Data Analysis

Acquired spectra were analysed using the Proteome Discoverer software suite (v2.3, Thermo Fisher Scientific) and the Mascot search engine (v2.6, Matrix Science^16^). Label free data were searched against SwissProt Human database. Data from the gel bands were searched against a custom “PHROG” database^13^ that includes 13 additional entries that correspond to mitochondrially encoded proteins for the *Xenopus* samples and the SwissProt mouse database for the mouse samples. TMT data was searched against the same custom “PHROG” database. False discovery rate (FDR) in peptide identification was set to a maximum of 5%. Peptide quantification data for the gel bands and the label free experiments were retrieved from the “Precursor ion area detector” node. The obtained values were used to calculate an estimation of protein amount with the top3 area. For the TMT data, peptides were quantified using the reporter ions intensities in MS3. Reporter ion intensities were adjusted to correct for the isotopic impurities of the different TMT reagents according to manufacturer specifications. For final analysis, values were transferred to Excel. For all experiments, identified proteins were selected as mitochondrial if they were found in MitoCarta 3.0^17^. MS3 spectra with abundance less than 100 or proteins with fewer than two unique peptides were excluded from the analysis. Each TMT channel was normalized to total mitochondrial protein abundance.

For human somatic cell samples, we analysed three dilutions: The 1X reference had similar protein loading to the primordial follicle sample (0,55 μg total protein); a 2-fold dilution (0,25 μg total protein) and a 5-fold dilution (0,1 μg total protein). In scatter plots (Fig 3g,h), we estimated differences in mitochondrial complex I protein abundance using the 2-fold somatic cell dilution, a conservative approach that compared primordial follicle samples (0,55 μg total protein) to somatic cells half their loading concentration (0,25 μg total protein), nevertheless observing similar levels of mitochondrial import machinery subunits TOMMs and TIMMs. 5-fold dilution somatic cell sample was useful for establishing detection limits, indeed, many complex I subunits absent in oocytes were detected with high confidence even in the 5-fold somatic cell dilution. In the heatmap (Fig 3i), we considered normalising our data using the mitochondrial loading controls citrate synthase and COX4I1 to estimate differences in protein abundance. The abundance of COX4I1 fell within the linear range of our proteomic methodology (R^2^=0.99 for COX4I1), in contrast to citrate synthase (R^2^=0.89 for CS) whose higher abundance led to measurement saturation at higher concentrations. Therefore, COX4I1 was chosen to normalize protein abundances in the heatmap representation.

The raw proteomics data have been deposited to the PRIDE repository^18^ with the dataset identifiers: PXD025366 (Quantitative proteomics), PXD025369 (Label-free) and PXD025371 (Gel band identification).

## Acknowledgements

We thank Anthony Hyman, Ben Lehner, Maya Schuldiner, Nicholas Stroustrup and members of the Böke lab for discussions and reading of the manuscript. We thank R. Barsacchi for assistance in the Seahorse assays. Graphical representations in figures 3a and S1f were created with BioRender. E.B. acknowledges support from MINECO’s Proyectos de Excelencia (BFU2017-89373-P), European Research Council Starting Grants (DORMANTOOCYTE - 759107) and support from the CRG Core Facilities for Advanced Light Microscopy, Histology, and Proteomics.

## Authors contributions

E.B. supervised the project. A.R-N. and E.B. designed the study, performed data analysis and wrote the manuscript. C.D.G and M.A.M-Z. provided human samples. A.R-N. performed all experiments except; ETC inhibition experiments performed by A.T-S., human primordial follicle isolations performed by A.R-N., J.M.D. and E.B., and MS sample preparation by E.B.

## Competing interests

The authors declare no competing interests.

---

**Figure S1.**
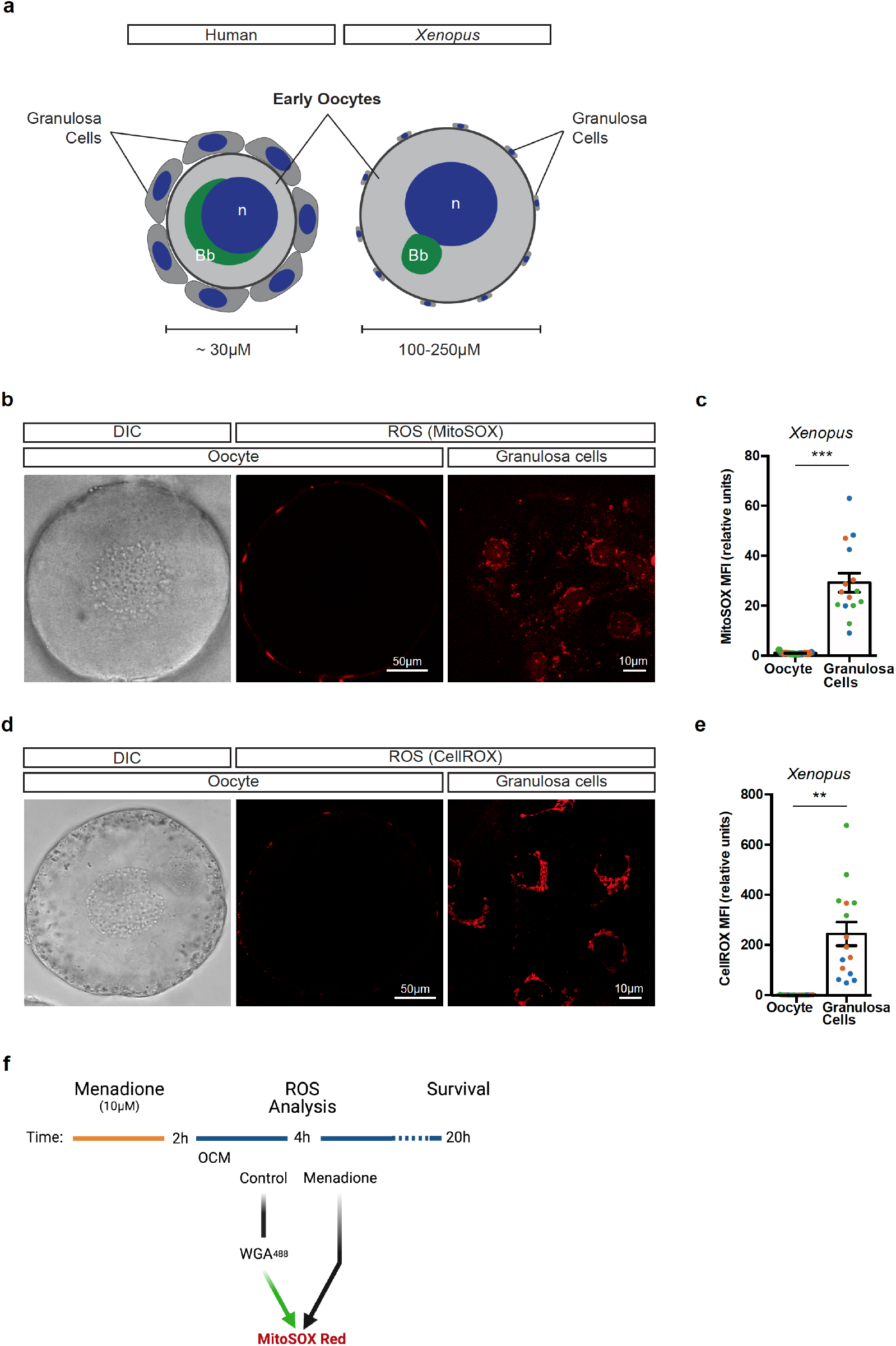
Undetectable levels of ROS in early oocytes. **a,** Schematic representation of human and *Xenopus* early oocytes with attached granulosa cells. Nuclei (n) are depicted in blue and Balbiani bodies (Bb) in green. Note that *Xenopus* early oocytes are huge, and their granulosa cells are only visible as little dots on the periphery of the oocyte in the same magnification. **b, d,** Live-cell imaging of *Xenopus* early (stage I) oocytes with attached granulosa cells with MitoSOX (**b**), and CellROX (**d**) to detect their ROS levels. Granulosa cells were imaged in the basal plane of the oocyte. Scale bars are as indicated in the figure. **c, e,** Quantification of MitoSOX (**c**) and CellROX (**e**) probes inside oocytes and in granulosa cells (n=3; biological replicates shown in colours). Data represent mean ± SEM. *** *p* = 4.298 × 10^-8^, ** *p* = 1.86 × 10^-5^ **f,** Experimental design for mild menadione treatment. Freshly isolated oocytes were treated with 10 μM menadione for 2 hours. ROS levels in oocytes were assessed 2 hours after menadione treatment in a fraction of treated oocytes. Untreated oocytes were labelled with Wheat Germ Agglutinin (WGA)^488^ to mark their plasma membrane and mixed with treated oocytes for MitoSOX labelling and imaging in the same plate. After menadione washout, oocytes were cultured in OCM overnight, and their survival was calculated.

**Figure S2.**
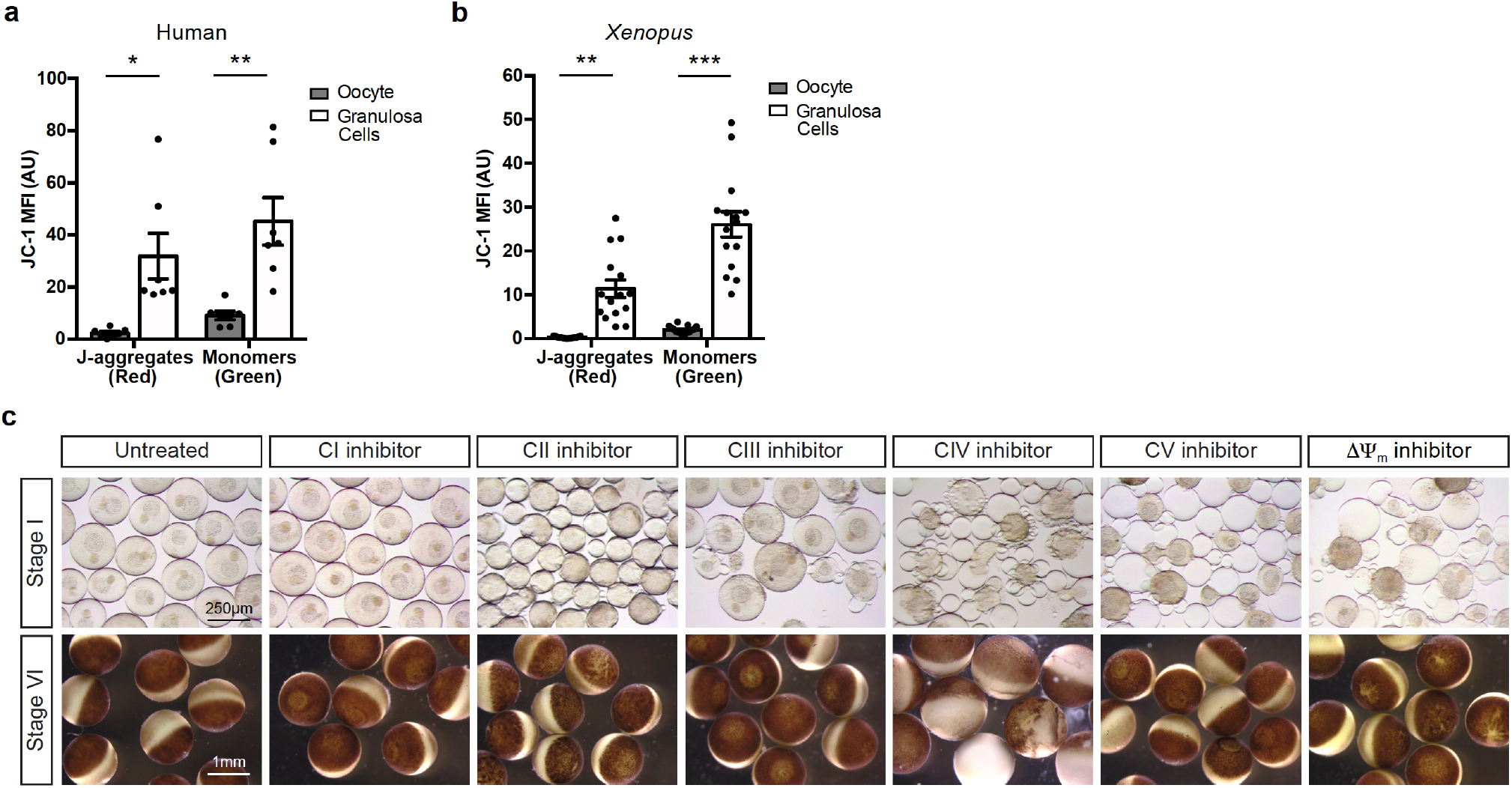
Oxidative phosphorylation is essential in early oocytes. **a,b,** Quantification of the mean fluorescence intensity (MFI) of JC-1aggregates (red) and monomers (green) in the oocyte and in the population of granulosa cells surrounding the equatorial plane of the oocyte for human (**a**) and *Xenopus* oocytes (**b**). We could not detect any red fluorescence in oocytes; an additional 2-hour incubation with JC-1 did not lead to detection of red fluorescence either inside the oocyte. Data represent mean ± SEM. * *p* = 0.005 and ** *p* = 0.002 in human, n=4; and ** *p* = 5.67 × 10^-6^ and *** *p* = 4.42 × 10^-9^, n=3 in *Xenopus*. **c,** Representative images of *Xenopus* oocytes after an overnight treatment with mitochondrial poisons (5 μM Rotenone, 50 mM malonic acid, 5 μM antimycin A, 50 mM KCN, 200 μM DCCD or 30 μM CCCP). Upper panel shows early (stage I) oocytes and bottom panel displays late (stage VI) oocytes. Cell death can be recognized in early (stage I) oocytes by loss of plasma membrane integrity and/or loss of nucleus; and in late (stage VI) oocytes by development of a mottling pattern in the pigmentation of the animal pole. Scale bars are as indicated in the figure.

**Figure S3.**
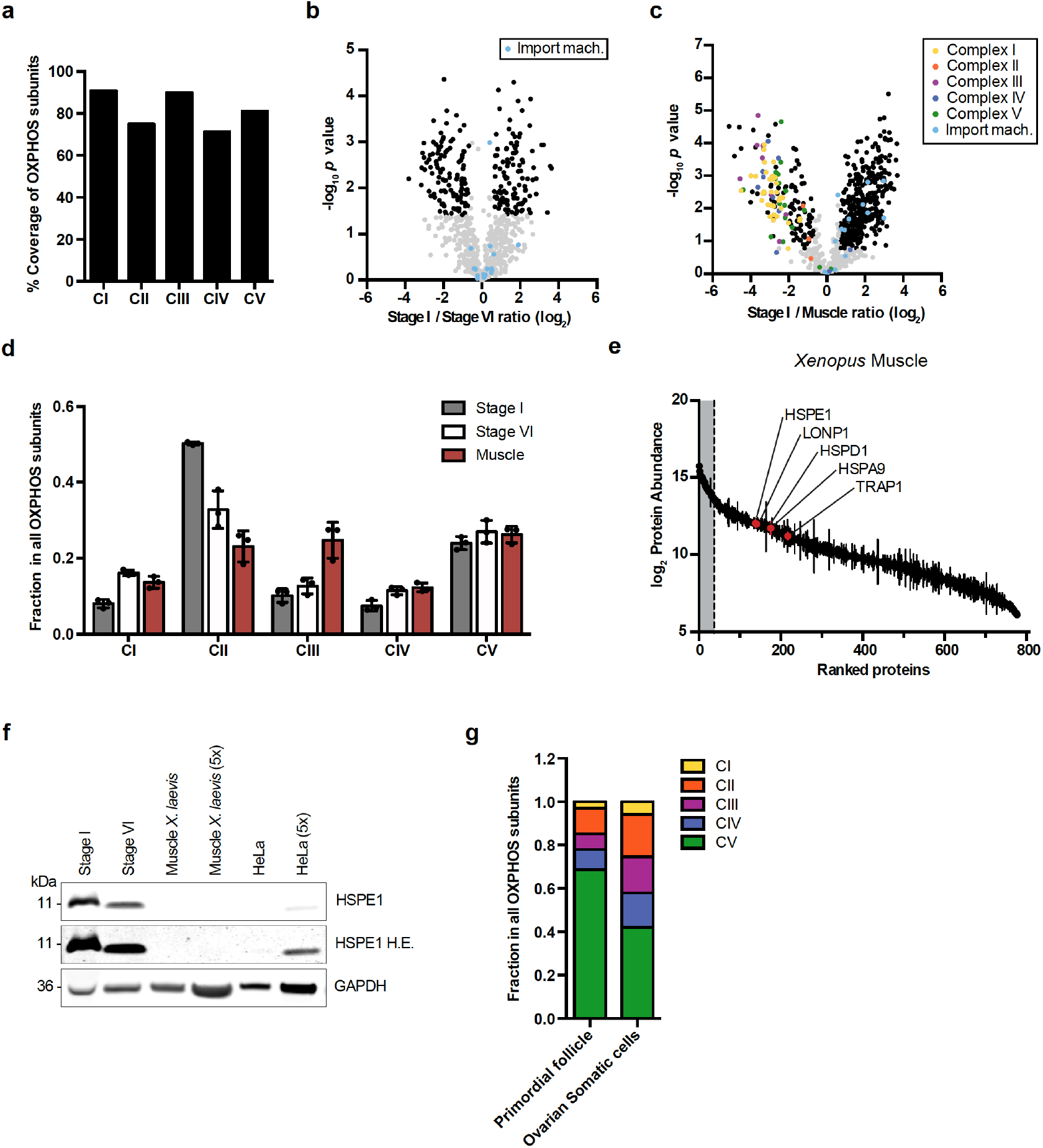
OXPHOS machinery is reduced in early oocytes. **a,** Percentage of proteins identified in the MultiNotch proteomics compared to all known subunits of OXPHOS machinery belonging to complexes I to V. **b,** Volcano plot showing *p* values versus fold changes of mitochondrial proteins between early (stage I) and late (stage VI) oocytes. Proteins significantly changing (q value <0.05, >1.5-fold change) are depicted in black. Subunits of mitochondrial import machinery (TIMs and TOMs) are highlighted in light blue. n=3 biological replicates. **c,** Volcano plot showing *p* values versus fold changes of mitochondrial proteins between early (stage I) oocytes and muscle. Subunits of OXPHOS and mitochondrial import machinery are highlighted in indicated colours. Other proteins significantly changing (q value <0.05, >1.5-fold change) are depicted in black. n=3. **d,** Fraction of the mean abundance of the indicated complexes among all of OXPHOS machinery in *Xenopus* oocytes and muscle tissue. Note that this analysis inevitably misses undetected subunits. Data are mean ± SEM for n=3 animals. **e,** Mitochondrial proteome in *Xenopus* muscle ranked by abundance. UPR^mt^ proteins are indicated in red. Data represent mean ± SEM from n=3 animals. **f,** Immunoblotting of HSPE1 in early (stage I), late (stage VI) oocytes, muscle and HeLa cells. GAPDH is used as a loading marker. H.E: High Exposure. **g,** Fraction of the mean abundance of indicated complexes among all of OXPHOS machinery in human follicles and ovarian somatic cells. CI - CV: Complex I - Complex V.

**Figure S4.**
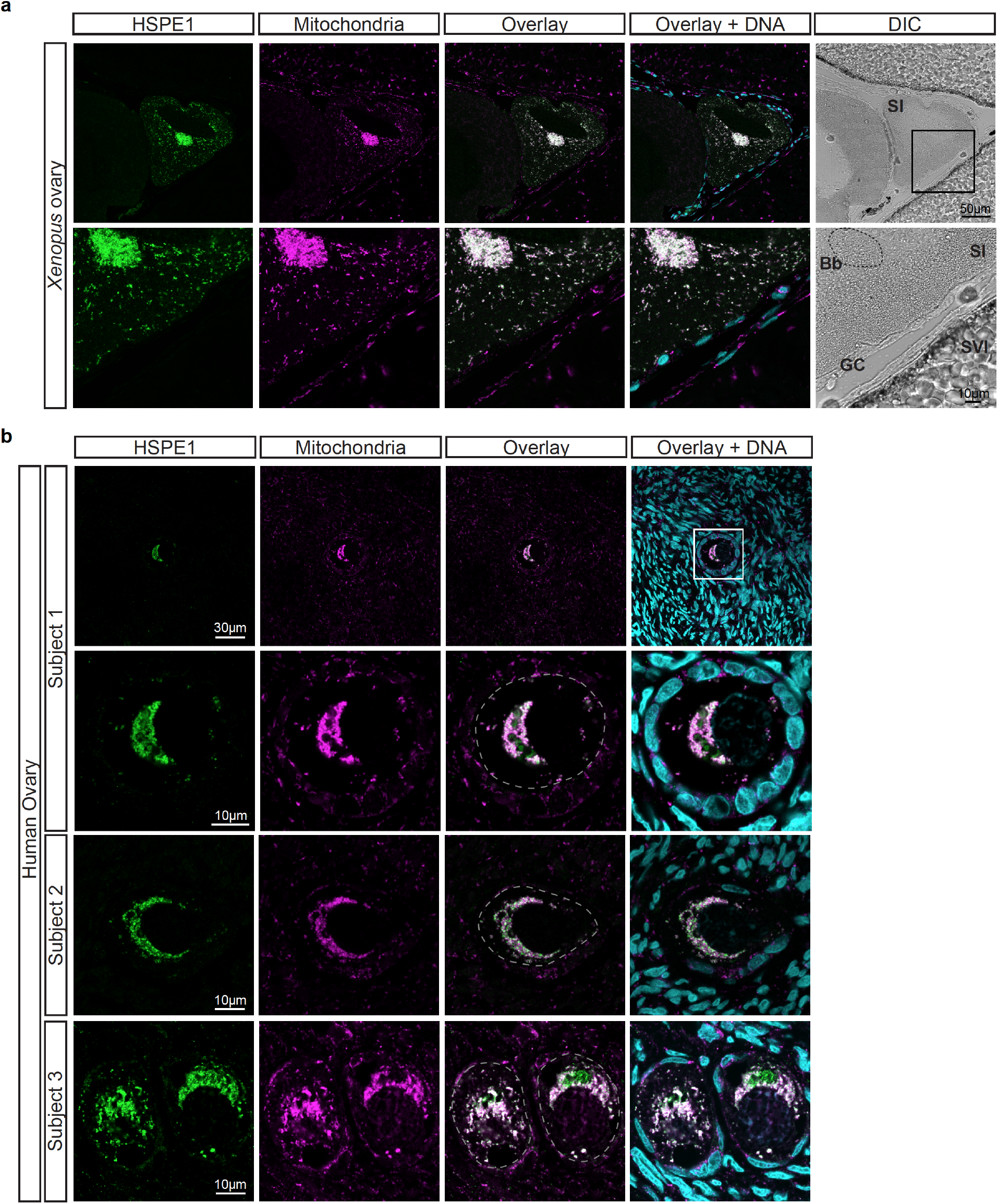
UPR^mt^ is constitutively active in early oocytes of human and *Xenopus*. **a, b,** Immunofluorescence of paraffin embedded sections of *Xenopus* (**a**) and human (**b**) ovaries with antibodies against HSPE1 (to monitor UPR^mt^) and ATP5A1 (to mark mitochondria). Hoechst was used to mark DNA. ATP5A1 was detected both in oocytes and somatic cells of the ovary. HSPE1, on the other hand, was so abundant in oocytes compared to surrounding cells that no signal could be detected from somatic cells without oversaturating the HSPE1 signal from early oocytes. SI: early (stage I) oocyte; SVI: late (stage VI) oocyte; GC: Granulosa cells; Bb: Balbiani body. Scale bars are as indicated in the figure.

**Figure S5.**
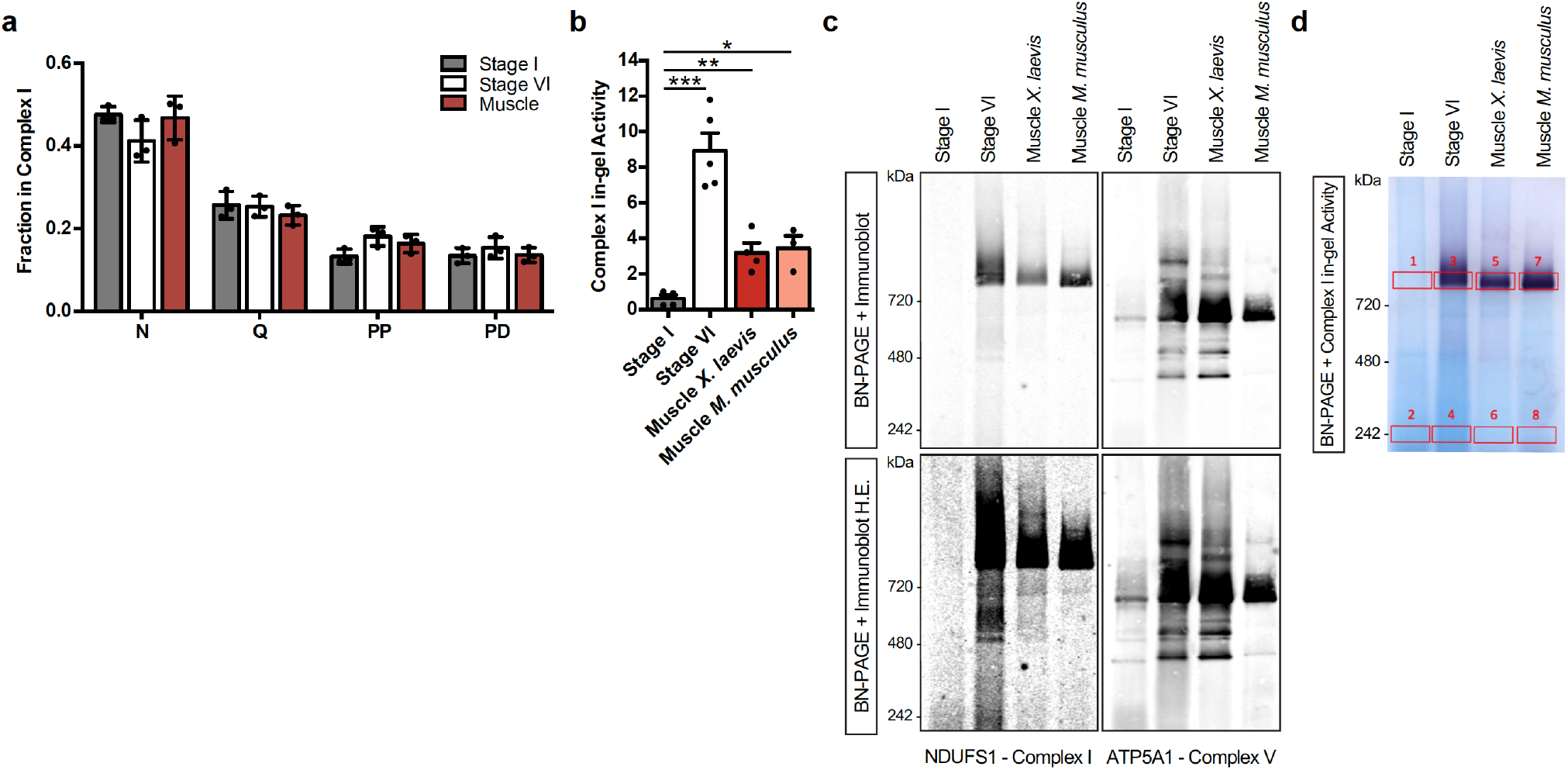
Complex I is not assembled in early oocytes. **a,** Fraction of the mean abundance of the indicated modules among all complex I modules in *Xenopus* oocytes and muscle tissue. Note that this analysis inevitably misses undetected subunits. Data are mean ± SEM for n=3 animals. **b,** Quantification of reduced NTB intensity in BN-PAGE gels after complex I in-gel activity assays. Data are mean ± SEM, n ≥ 3, * *p* = 0.0209, ** *p* = 0.017, *** *p* < 0.0001. **c,** DDM solubilized mitochondrial fractions were resolved by BN-PAGE followed by immunoblotting using antibodies against NDUFS1 and ATP5A1 to detect complexes I and V, respectively. Lower panels show high exposure (H.E.) blots. **d,** Representation of excision sites for mass spectrometric identification of complex I and complex II subunits in Fig 4c.

## Notes

### Competing Interest Statement

The authors have declared no competing interest.

